# Single nucleus RNA-sequencing reveals transcriptional synchrony across different relationships

**DOI:** 10.1101/2024.03.27.587112

**Authors:** Liza E. Brusman, Julie M. Sadino, Allison C. Fultz, Michael A. Kelberman, Robin D. Dowell, Mary A. Allen, Zoe R. Donaldson

**Affiliations:** Department of Molecular, Cellular, and Developmental Biology, University of Colorado Boulder; Boulder, CO 80309 USA; Department of Psychology and Neuroscience, University of Colorado Boulder; Boulder, CO, 80309 USA; Biofrontiers Institute, University of Colorado Boulder; Boulder, CO, 80309 USA

## Abstract

As relationships mature, partners share common goals, improve their ability to work together, and experience coordinated emotions. However, the neural underpinnings responsible for this unique, pair-specific experience remain largely unexplored. Here, we used single nucleus RNA-sequencing to examine the transcriptional landscape of the nucleus accumbens (NAc) in socially monogamous prairie voles in peer or mating-based relationships. We show that, regardless of pairing type, prairie voles exhibit transcriptional synchrony with a partner. Further, we identify genes expressed in oligodendrocyte progenitor cells that are synchronized between partners, correlated with dyadic behavior, and sensitive to partner separation. Together, our data indicate that the pair-specific social environment profoundly shapes transcription in the NAc. This provides a potential biological mechanism by which shared social experience reinforces and strengthens relationships.

## Main Text

Social bonds are shaped through the continuous integration of shared experiences, resulting in a mutual understanding that is critical for successfully navigating real-time interactions and achieving long-term collaboration. Yet how a pair is able to coordinate their behavior has been a long-standing question in social neuroscience. Non-invasive human research has repeatedly highlighted interbrain neural synchrony – the alignment of oscillatory neural activity between individuals – as an emergent property of social interaction (*1*, *2*). Interbrain synchrony scales with relational closeness, and is associated with enhanced empathy, liking, rapport, and prosocial behavior (*3–9*). In animal models, neural synchrony has been observed during acute social interaction (*9*, *10*), decision making in social dominance tests (*8*), and cooperation tasks (*11*).

While neural synchrony may initially arise from real-time interpretation of the same stimuli, its strengthening between individuals in a close relationship likely reflects aligned cognitive processing, even in the absence of shared cues (*12*). In the absence of shared cues, neural synchrony and organized intra-pair behavior must rely on pre-existing, common biological substrates that prime neurons to fire synchronously. This underlying state is likely achieved through shared prior experiences that have altered cell states in the same way across individuals.

Thus, we would expect this shared biological state to emerge as a product of social experience with another individual. In prairie voles, correlation in *cFos* expression – a molecular marker of neuronal activity – emerges between partners in some brain regions within the first 24 hours of pairing (*13*). Further, in pairs of fighting betta fish, brain-wide gene transcription becomes more correlated between partners, but not between non-partners, as the duration of fighting increases (*14*).

In the present study, we investigated if a pair-specific global transcriptional state exists in the context of different types of social bonds. To determine whether a pair-specific signature exists in the nucleus accumbens (NAc), a ventral striatal region extensively implicated in social bonds and important for reward, learning, and action selection, we performed single nucleus RNA-sequencing (snRNA-seq) of the NAc in prairie voles. In contrast to the vast majority of single cell (sc)/snRNA-seq studies to date, rather than pooling samples, we sequenced the NAc from each vole individually. This provided a unique opportunity to assay gene-behavior correlates at the individual level and query intra-pair transcriptional synchrony. We leveraged this approach to compare within-pair transcriptional synchrony to across-pair synchrony as a powerful internal control. Together, this experimental design interrogates intra-pair similarity that is not simply a consequence of a general housing environment, sex, or bonding state in voles more broadly.

Prairie voles form selective affiliative relationships with peers or with a mate, akin to human friendships and romantic relationships, respectively (*15*, *16*). We identified distinct gene modules that underpin individual variation in bond-related behaviors in different sexes/relationship types. Using machine learning and more traditional analytical approaches, we further found that voles were more transcriptionally similar to their relationship partner than other voles regardless of sex or pairing type, which we refer to as transcriptional synchrony. We then show that synchronized gene expression is correlated with pairwise interaction behavior and highly sensitive to partner loss. Our data are the first to identify a potential cell-level molecular mechanism underlying organized intra-pair patterns of neural activity (e.g. synchrony) and behavior, providing novel insights into how pair-specific behaviors may be reflected at a cellular level.

### Prairie vole NAc transcriptional landscape across relationship types

We used snRNA-seq to identify cell type-specific NAc expression patterns that support long-term social bonds using prairie voles. As in humans, peer and mate relationships in this species share many behavioral features, with reproductive behaviors predominantly exhibited within mate relationships. To examine transcriptional differences and similarities across different types of relationships, we paired individual prairie voles with either a novel same-sex (peer-paired) or opposite-sex (mate-paired) partner.

To map the transcriptional landscape of the prairie vole NAc, we collected NAc tissue 16 days post-pairing (1 day post-behavior testing). 39 of 40 tissue samples passed quality control and were merged to create a combined dataset of 142,488 single nuclei (Table S1; 8 peer-paired females, 10 peer-paired males, 11 mate-paired females, and 10 mate-paired males). In pair-wise analyses, the unpaired animal (one female from a mate pair) was excluded. We identified 15 transcriptionally-defined cell type clusters in the NAc based on known markers of cell types in this region (Fig 1A, B). Medium spiny neurons (MSNs), the primary neuronal cell type within the NAc, were identified based on their combined expression of the dopamine receptors *Drd1a* and *Drd2*, the opioid receptor *Oprm1*, and opioid ligands *Pdyn* and *Penk*. This resulted in five MSN clusters we termed MSN-Drd1Pdyn, MSN-Drd1PdynOprm1, MSN-Drd1Penk, MSN-Drd2Penk, and MSN-Drd2NoPenk. Five interneuron clusters were identified based on their expression of only *Gad1/Syt* (Int-GABAergic), *Igfbpl1/Dlx2* (Int-Dlx2Immature), *Sst/Npy* (Int-SstNpy), *Vip/Kit* (Int-Pvalb), and *Chat* (Int-Cholinergic). Glia were identified based on their expression of *Mog* (MatureOligos), *Olig2/Pdgfra* (oligodendrocyte progenitor cells, or OPCs), *Gja1* (Astrocytes), *Aif1* (Microglia), and *Vim* (RadialGlia-LikeCells). These clusters are largely consistent with those identified in snRNA-seq datasets from other species including mice (*17*), rats (*18*), and humans (*19*), suggesting overall cell type conservation in the NAc, congruent with a conserved role for this brain region in learning, reward and social behavior across species (*20*).

**Figure 1.**
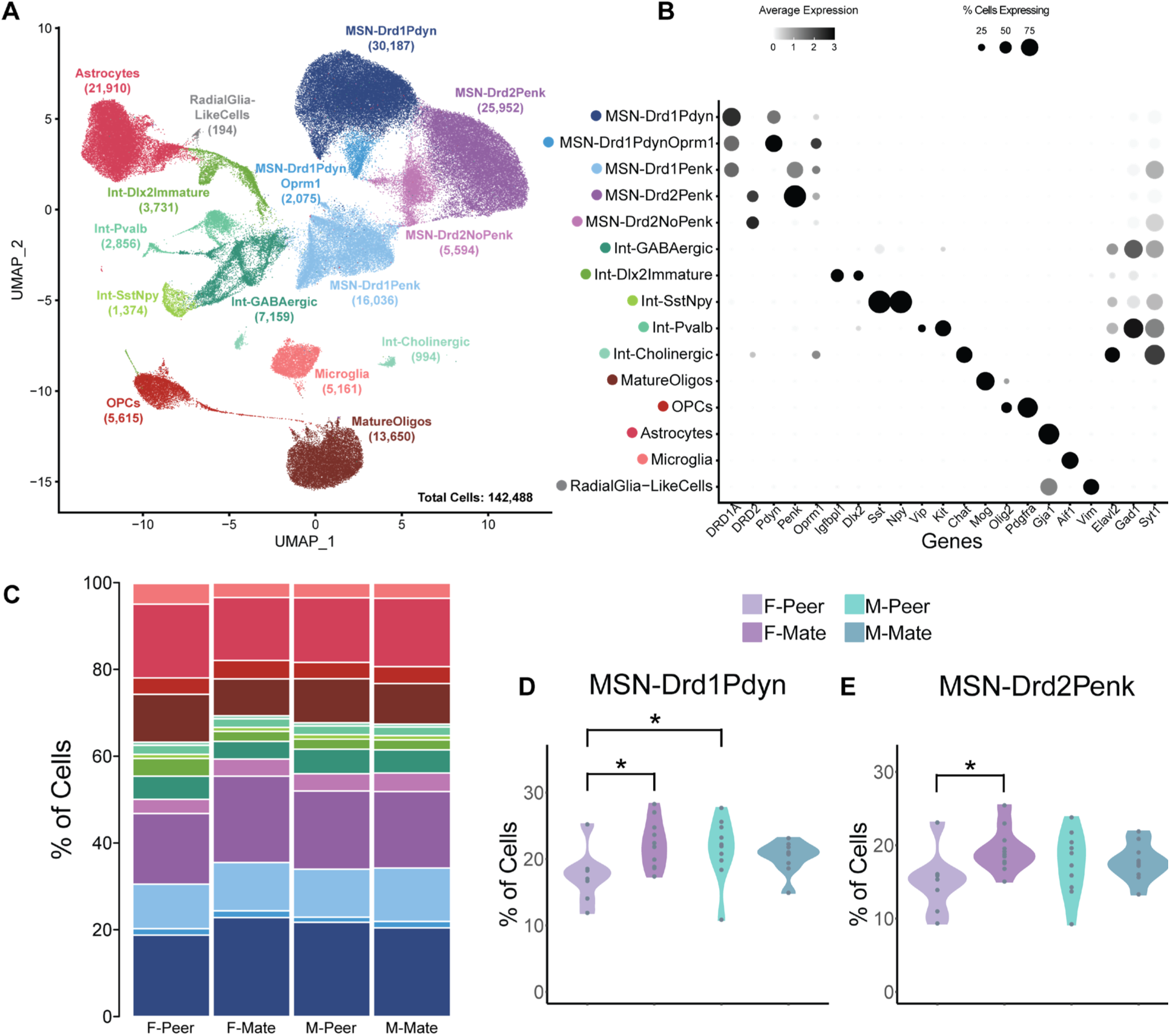
Prairie vole NAc transcriptional landscape across relationship types. **A.** UMAP clustering of NAc cells revealed 15 clusters. **B.** Dotplot indicating expression of marker genes. **C.** Stacked bar plot of cell type proportions calculated per experimental group (colors refer to cell groups in A, B). **D, E.** Cell type proportions differ between groups in MSN-Drd1Pdyn and MSN-Drd2Penk clusters. * p < 0.05.

Having identified the major cell classes that comprise the NAc, we next asked whether cell type proportions differed by sex or pairing condition. For most clusters (13/15), the proportion of cells in each cluster did not differ between sexes or pairing types (Fig S1). However, mate-paired females had a higher proportion of MSN-Drd1Pdyn and MSN-Drd2Penk neurons than peer-paired females (GLM with estimated marginal means, FDR corrected: MSN-Drd1Pdyn: T = 2.93, p = 0.024; for MSN-Drd2Penk: T = 2.80, p = 0.033), and peer-paired males had higher proportion of MSN-Drd1Pdyn neurons than peer-paired females (T = 2.44, p = 0.041; Fig 1C, D, E). This proportional shift is likely due to shifts in neuronal gene expression rather than neurogenesis as the overall number of MSNs does not differ across groups. This is consistent with prior work indicating that dopamine D1-class receptors and the kappa opioid receptor ligand *Pdyn* are upregulated in the NAc upon bonding in prairie voles (*21*, *22*). Such changes may reflect the onset of selective aggression (mate guarding) in mate relationships (*21*, *22*).

To identify transcriptional correlates of pair bonding behavior, we performed partner preference tests (PPTs) after 14 days of pairing on the same 20 vole pairs (4 female/female or FF, 5 male/male or MM, 11 female/male or FM) that were sequenced (Fig 2A, B). After excluding the one mate-paired male whose sequencing did not pass quality control, 38 out of 39 voles displayed a partner preference (>50% time huddling with partner vs. novel; Fig 2C, D). There were no group differences in partner preference metrics based on sex or pairing type (Fig 2C, D). These results are consistent with prior reports (*15*, *16*, *23*) and provide a platform to identify the unique and shared features of different types of social bonds.

**Figure 2.**
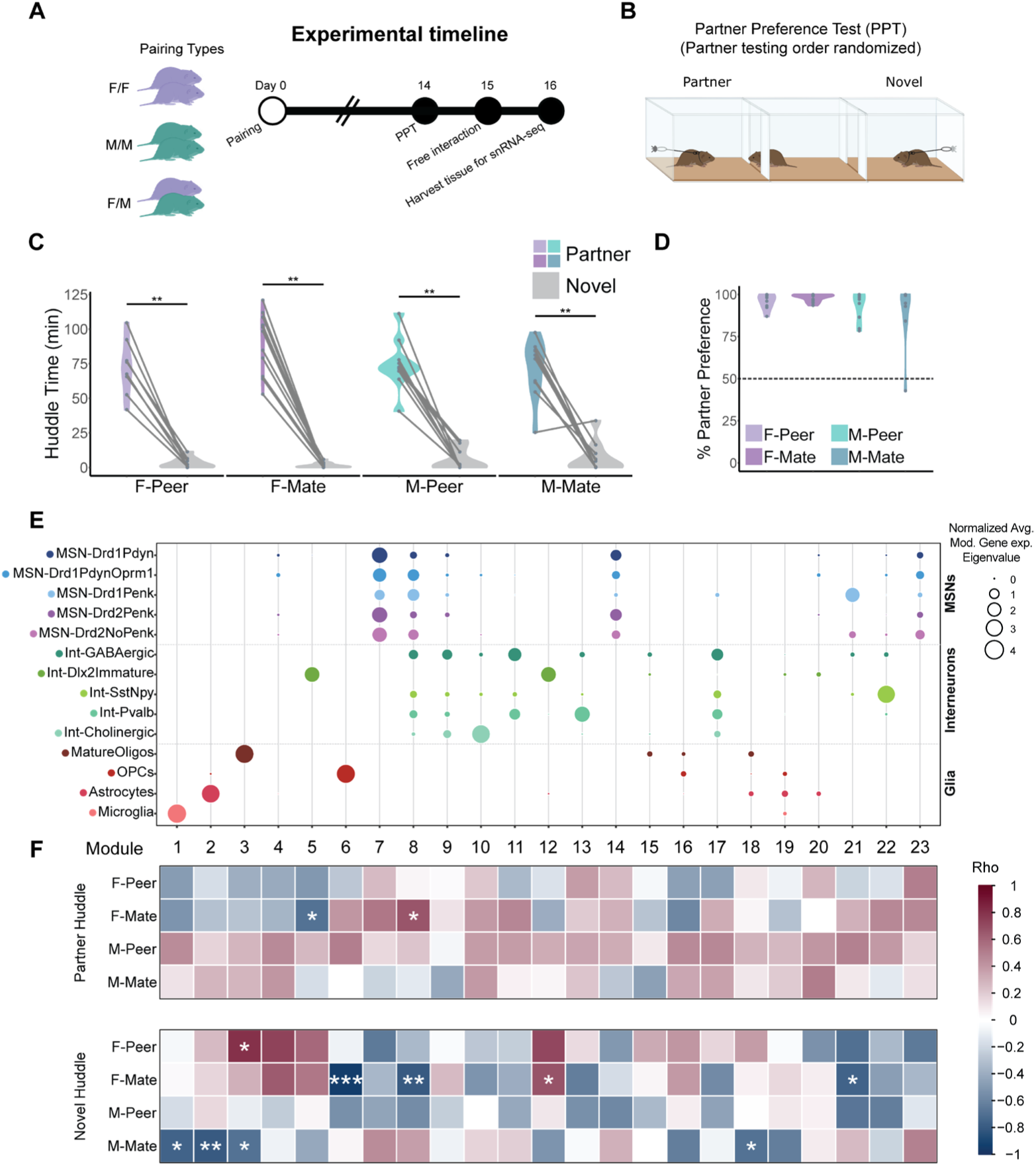
Gene module expression associations with behavior in different relationship types. **A.** Experimental timeline. **B.** Schematic of partner preference test (PPT). **C.** Huddle time in the partner preference test. All groups huddle significantly more with the partner than the novel vole. There are no group differences in partner huddle or novel huddle time. **D.** Percent partner preference calculated as [partner huddle]/[partner huddle + novel huddle]x100%. All groups show a partner preference (mean % partner huddle > 50%) and there are no group differences. **E.** The top 3,000 variable genes identified by Hotspot were grouped into 23 co-expression modules. The size of the circle indicates the average module eigenvalue (calculated per cell) for cells in each UMAP cluster. **F.** Module expression correlation with PPT behaviors (top: partner huddle, bottom: novel huddle). The color of the box indicates Spearman’s Rho. Significant correlations (corrected via bootstrapping) are indicated with asterisks. * p < 0.05, ** p < 0.01, *** p < 0.001.

### Gene expression patterns underlying different relationship types and individual differences in relationship behaviors

We next examined gene-behavior relationships using Hotspot, a cluster-agnostic computational algorithm that groups genes into modules based on similar expression patterns across cells (*24*, *25*) (Table S2). Seurat and Hotspot differ in their analytical purposes, with the first designed to cluster cell types and the latter to identify modules of genes with similar expression patterns. Thus, the top 3,000 variable genes identified by our cell clustering algorithm (Seurat) only partially overlapped (1,954/3,000, or 65%) with the top 3,000 variable genes identified by Hotspot.

Using Hotspot, we identified 23 gene modules (31-415 genes/module) that are informative to, but not constrained by, cell type clusters (Fig 2E). For instance, modules 7 and 14 consist of genes expressed across the MSN cell clusters, enabling us to query shared gene expression differences across these cell types as they relate to behavior. Surprisingly, we found that no modules differed at the group level (Fig S2), suggesting that individual variation in gene expression supersedes sex and pairing type differences and that gene expression is largely shared across different types of bonds.

To better understand what factors may be associated with gene expression variability between animals, we used the Hotspot gene modules to interrogate gene-behavior relationships at the individual level. We calculated the average (mean(log(counts))) expression of genes for each animal in each module and compared module expression to behavior in the PPT (Spearman correlation with significance threshold determined by bootstrapping). We found that different gene modules correlated with PPT behavior in different sex/pairing type groups (Fig 2F). This suggests that different gene networks regulate the same behaviors in different social relationships. Although the specific modules that exhibit gene-behavior correlations differ between groups, 3 of our 4 groups (F-Peer, F-Mate, M-Mate) show correlations between novel-directed behavior and gene expression in oligodendrocytes (Module-3 in MatureOligos and Module-6 in OPCs; Fig 2E, F). This is consistent with prior work from our lab implicating oligodendrocytes in pair bonding (*23*) and with a role for oligodendrogenesis and myelination in complex learning (*26–28*). Further, most gene-behavior correlations were in novel-directed rather than partner-directed behavior, suggesting that these gene programs play a role in the dissociation of pro- and anti-social behaviors that comprise behavioral selectivity within relationships.

### Transcriptional synchrony within pairs

We next tested the hypothesis that transcription would be more similar between partners than between non-partners (Fig 3A, B). To leverage the unique advantages of our individual animal and cellular-resolution dataset, we developed a transcriptional decoder, which uses the multidimensional transcriptional profile (expression of the 3,000 variable genes from Seurat) to predict the vole from which a cell originated. Specifically, we used support vector machines (SVMs) as multi-class classifiers. SVMs work by finding differences between samples and grouping them into distinct categories. Here, we used this principle to examine similarity between cells from different animals; any (mis)classification of an individual cell reflects the animal whose transcriptional profile that cell is most alike.

**Figure 3.**
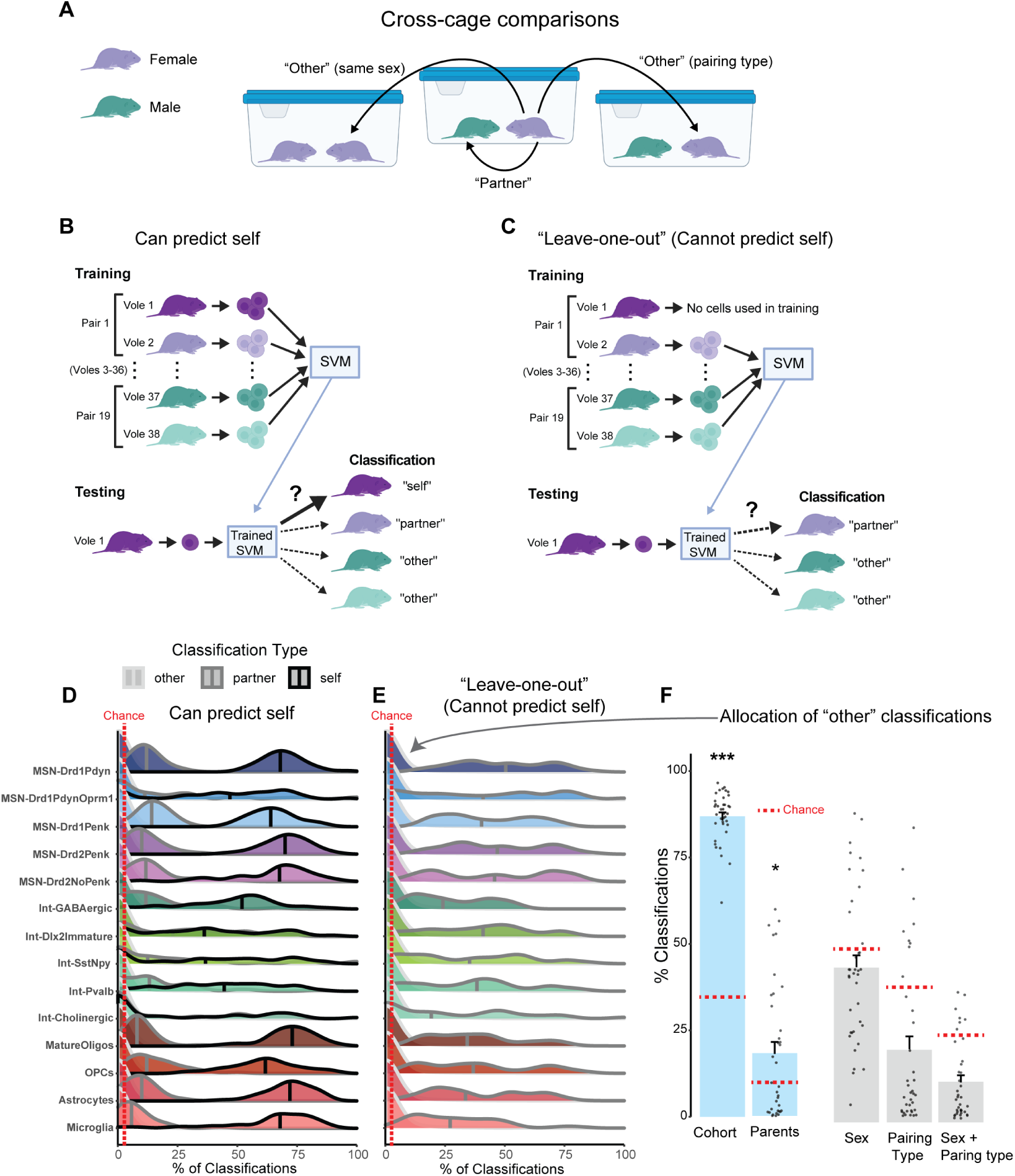
SVM-based decoding of cellular transcriptional profiles reveals similarity between partners. **A.** Schematic of intra- and inter-cage comparisons (“partner” and “other”, respectively). **B.** Schematic of how SVMs were trained to predict the identity of the originating vole for each cell. Correct predictions are denoted as “self”. **C.** Schematic of how SVMs were trained to predict the most similar transcriptional profile in a hold-one-out design where assignment of self was not possible. **D.** The SVM predominantly classified cells correctly (self, black outline), with the most common misclassification assigned to their partner (dark grey outline), followed by any other animal (light grey outline). **E.** SVMs were iteratively trained for each cluster by holding one animal out from the training set, and then testing the SVM on the held-out animal. The held-out animal was most frequently classified as their partner (dark grey outline) rather than any other animal (light grey outline). **F.** Allocation of “other” classifications from the hold-one-out SVMs (**E**). The y-axis indicates the percentage of “other” cells for each animal belonging to animals from the same cohort, same parents, same sex, same pairing type or sex-by-pairing type. The red dashed line denotes the expected percentage of classifications for animals in the same category for each metric (chance). * p < 0.05, ** p < 0.01, *** p < 0.001

We trained our transcriptional decoder in a cell cluster-specific manner using 75% of our dataset (up to 150 cells/cluster/vole), reserving the remaining 25% (up to 50 cells/cluster/vole) as the test dataset. The chance likelihood that the classifier assigns a cell from the dataset to the correct vole is only 2.6% (i.e. 1 out of the 38 voles in our dataset). We separated the resulting cell classifications from the SVM into three categories: cells originated from the correct animal (“self”), the correct animal’s partner (“partner”) or any other animal (“other”). Our classification strategy and experimental design enables important cross-pair comparisons that serve as robust internal controls (Fig 3A, B). Cross-pair comparisons are more informative than comparing to animals housed in isolation, as isolation has profound effects on neural gene expression (*23*, *29*), and the lack of a partner makes it impossible to test for pairwise synchrony. Rather, reliable decoding of identity when comparing animals housed *in the same type of relationship* provides much stronger evidence of intra-pair transcriptional synchrony.

Supporting the validity of our machine learning approach, we found that, in every cluster, our SVMs are most likely to classify cells correctly (e.g. “self”, Fig 3D). The mean percentage of self-classifications ranged from 24.9% to 72.2% across clusters. When cells were misclassified by the SVM, they were much more likely to be classified as the partner animal (“partner”) than any other non-partner animal (“other”) (Fig 3D, p < 10^−5^ for all partner vs. other comparisons). This was not sensitive to sex or relationship type (Figs S3, S4). Mean partner classifications ranged from 7.4% to 25.3% across clusters, still significantly greater than chance (2.6%) albeit lower than self. Moreover, this was not due to cohort effects (Fig S5), providing further evidence that partners are more transcriptionally similar than non-partners.

To further test the hypothesis that partners show transcriptional synchrony, we employed a “leave-one-out” version of our decoder. We tasked an SVM with classifying cells when the correct choice (“self”) was not available, forcing the classifier to assign the cell to an animal other than self (Fig 3C). Strikingly, when correct self-assignment was not available, the SVMs were most likely to classify the cells as originating from the partner animal (Fig 3E, p < 10^−16^ for partner vs. other in all clusters). The mean percentage of partner classifications ranged from 27.3% to 48.5% across clusters. While this was lower than accurate self-classification, it dramatically exceeded classifications for “other” and random chance, which would predict a partner classification only 2.7% of the time (1 of 37 voles) (Fig 3E, p < 10^−5^ for chance vs. partner in all clusters).

While the results of our decoder indicate that partners are more transcriptionally similar than non-partners, this does not rule out a potential role for other factors in driving transcriptional similarity to some extent. We collected behavior and tissue in three cohorts of animals, run in successive waves constrained by the number of behavioral tests that could be run in parallel. All nuclear isolation and sequencing randomly incorporated animals across cohorts. Using the “leave-one-out” classifiers that could not predict self (Fig 3C, E), we asked if the cells classified as “other” were from the same experimental behavioral cohort, the same group (F-Peer, F-Mate, M-Peer, M-Mate), the same pairing type (FF, FM, MM), the same parents (i.e. siblings), and the same sex. We found that, compared to the expected percentage of classifications, behavioral cohort has the largest effect on cell classification with cells being ∼2.5 times more likely to be assigned to the same cohort than would be expected by chance (Fig 3F, p < 10^−15^, T = 37.08). This indicates that animals from the same behavioral cohort, which existed at the same time in our colony and underwent the experimental timecourse together, are more transcriptionally similar to each other than to animals from other cohorts. Additionally, animals were more likely to be classified as sibling animals (same parents) than non-siblings, although with a much smaller effect size (∼0.75X greater than chance) (Fig 3F, p = 0.013, T = 2.62). This suggests that there is some genetic contribution to transcriptional similarity albeit less than shared cohort effects. Cells were classified approximately equally as originating from an animal of the same or opposite sex (Fig 3F), reflecting a potential lack of overall sex differences in the NAc in prairie voles. The other factors we investigated also do not exceed chance probability.

In sum, these data demonstrate that SVMs are a powerful tool for inference of cell identity, one that we have employed to discover factors important for driving transcriptional similarity across individuals. Partner is the predominant factor for explaining transcriptional similarity between two voles, although we also identified a smaller role for cohort and genetic relatedness. However, even classifications for other cohort members did not exceed classifications for a partner (Fig S5). This supports our hypothesis that it is *social* environment and an animal’s partner specifically that is the largest contributor to transcriptional synchrony.

### Genes responsible for transcriptional synchrony

To identify genes that play a role in transcriptional synchrony, we examined pairwise relationships in Hotspot module gene expression. We found that, regardless of pairing type, two Hotspot modules, Module-6 (expressed in OPCs) and Module-11 (expressed in interneurons), exhibited pairwise expression similarity via multiple metrics. First, Module-6 and Module-11 average gene expression is correlated between partners (Module-6: Rho = 0.61, p = 0.0057; Module-11: Rho = 0.59, p = 0.0075, Fig 4A, C). Second, using the expression of all genes within each module to calculate the Euclidean distance between partners, we found that for these two modules, partners were closer together in Euclidean space than non-partners (Wilcoxon rank sum tests: Module-6: W = 9781, p = 0.00017; Module-11: W = 8803, p = 0.0083, Fig 4E, F, G). Finally, when we ranked the animals by their module gene expression and calculated the rank-based distance between partners, we found that partners were closer together in the ranking than would be expected by chance (One-sample t-tests vs. null (expected) value: Module-6: T = −3.46, p = 0.0028; Module-11: T = −2.73, p = 0.014, Fig 4H, I). In all instances, there was no difference in the degree of pairwise similarity (via correlation, Euclidean gene expression distance, or rank) based on relationship type (Figs S6, S7, S8).

**Figure 4.**
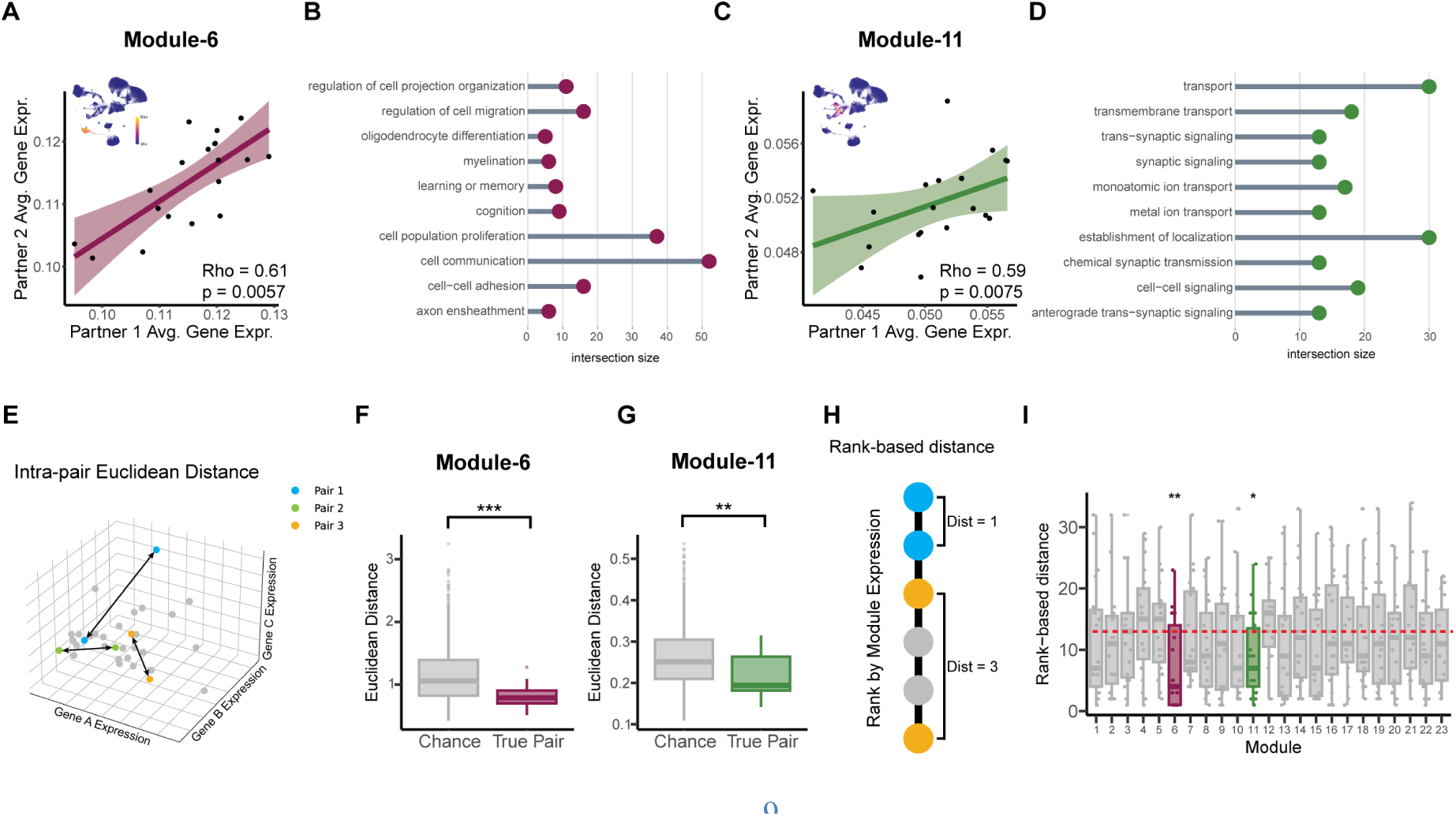
Module correlation between partners and with dyadic behavior. **A, C.** Spearman correlation between partners for average module gene expression of Module-6 (**A**) and Module-11 (**C**). **B, D.** Selected GO terms for Modules-6 (**B**) and −11 (**D**), prioritized based on relevance for neural/glial function. Intersection size indicates number of genes in module overlapping with genes in that term. **E.** Diagram of intra-pair Euclidean distance calculation based on counts for genes in each module. **F, G.** Euclidean distance between partners (True Pair) is less than between all non-true pair distances (Chance) for Modules-6 (**F**) and −11 (**G**). **H.** Diagram of pairwise rank-based distance calculation based on module gene expression. **I.** Rank-based distance between partners for each module is less than chance for Modules-6 and −11 (line at y=13 represents average chance distance). * p < 0.05, ** p < 0.01, *** p < 0.001.

GO terms for Module-6, include terms related to oligodendrocyte differentiation and myelination, and GO terms for Module-11 include terms related to synaptic signaling and ion transport (Fig 4B, D). The former suggests that oligodendrocyte maturation, subsequent myelination by mature oligodendrocytes, and/or synaptic pruning by oligodendrocytes could be contributing to similarity in population-level neural activity in partners. In contrast, the latter may act on faster timescales consistent with a role for interneurons in shaping population- and ensemble-level activity within the striatum (*30–32*). Together, these data indicate that specific gene networks in OPCs and interneurons are synchronized between partners and that oligodendrocyte maturation and interneuron signaling may be drivers of pair-wise transcriptional similarity regardless of pairing type.

### Transcriptional synchrony and pairwise interaction behavior

Given that partners exhibited correlated expression of Modules-6 and −11, we asked whether expression of these modules predicted pairwise behavior. On day 15 after pairing, we performed three-hour free interaction tests as a metric of dyadic behavior (Fig 5A). Total interaction time did not differ between pairing types (Fig 5B). Unlike in the PPT, where the tethered animals lack some amount of agency and choice, during free interaction both members are allowed to initiate or reject interaction. This better reflects ethologically relevant social behaviors and captures the complexity of a two-body system. Further supporting that these two tests measure different facets of social behavior, we replicated prior findings that partner huddle in the PPT is not predictive of levels of dyadic interaction in the free interaction test (Rho = 0.04, p = 0.8061; Fig 5C) (*33*).

**Figure 5.**
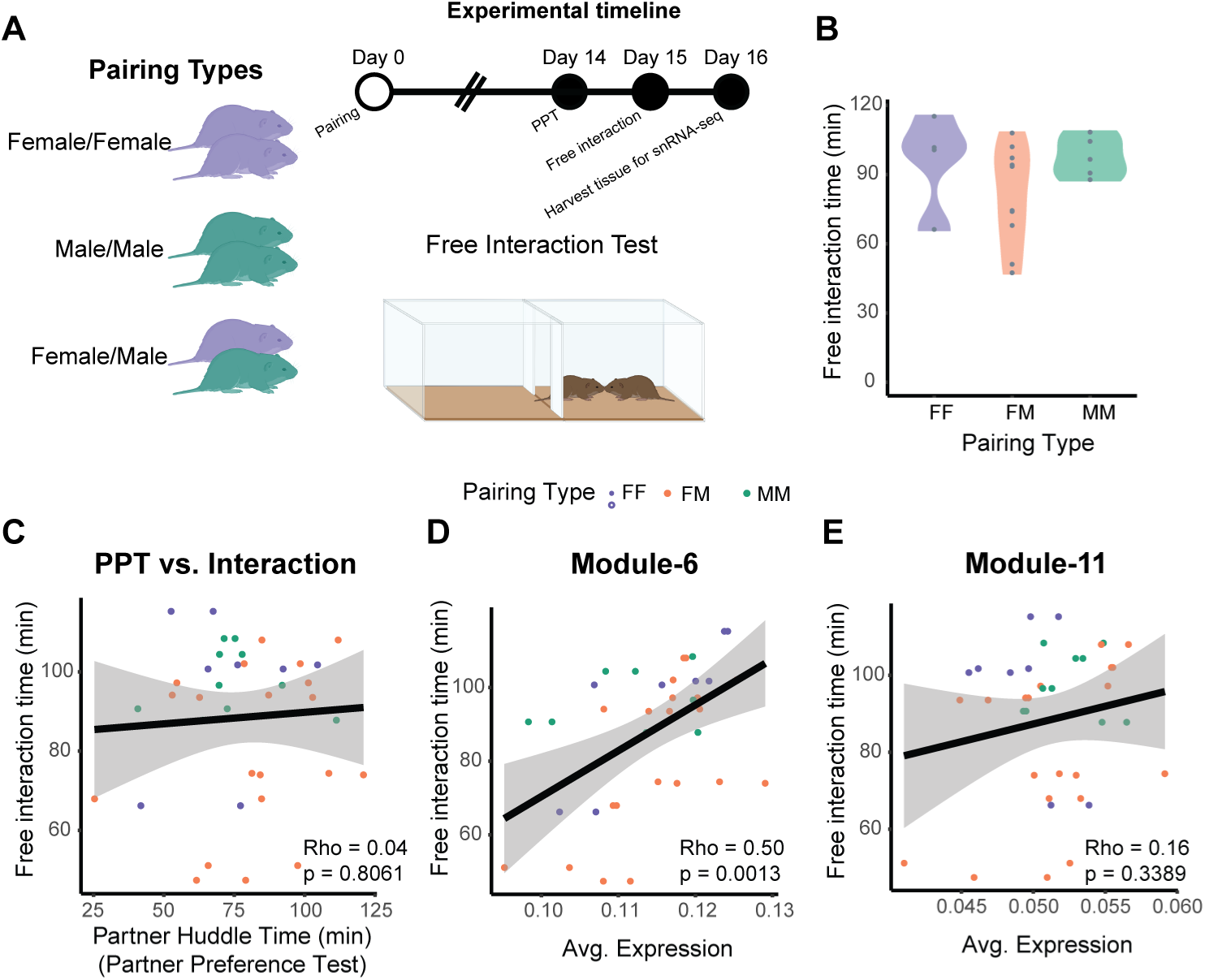
Synchronized Module-6 expression is associated with pairwise behavior. **A.** Experimental timeline and schematic for free interaction test. A video of the interaction test is also available in the online version. **B.** Total time interacting with a partner in the free interaction test does not differ between groups. **C.** Partner-directed huddle in the PPT does not predict pairwise interaction. **D, E.** Average gene expression for Module-6 (**D**) but not Module-11 (**E**) correlates with interaction time in the free interaction test.

We next found that synchronized gene Module-6 was correlated with pairwise interaction (Module-6: Rho = 0.50, p = 0.0013, Module-11: Rho = 0.16, p = 0.34, Fig 5D, E). Thus, expression of a synchronized gene module in OPCs is associated with pairwise behavior regardless of bond type. This supports our hypothesis that the pair-specific bond itself – rather than the type of relationship – is a critical factor in transcriptional and behavioral synchrony.

### Synchronized genes are sensitive to bond loss

Module-6 genes are correlated between partners and with pairwise dyadic behavior, leading us to hypothesize that these genes would be sensitive to manipulation of the bond itself and social context more generally. To test this hypothesis, we used data previously generated in our lab in which male voles were paired with either a peer or mate partner for 2 weeks (equivalent to our snRNA-seq timeline) or paired for 2 weeks and then separated from their partner for an additional 4 weeks (*23*). This timecourse is validated by prior work showing partial erosion of bonds (*34*) and potential for re-bonding at this timepoint (*35*). We then performed tissue-level RNA-seq of the NAc from these paired and separated animals. We identified 82% (117/143) of our Module-6 genes in our tissue-level RNA-seq data. Upon clustering the mean z-scored expression values for each group, we found that our genes grouped into 3 major clusters: one that has higher expression in separated vs. paired animals for both peer and mate bonds, one that has lower expression in separated vs. paired animals for both peer and mate bonds, and one cluster that does not show consistent expression changes between peer- and mate-bonded animals (Fig 6A, B, C). 71% (83/117) of our Module-6 genes show consistent differences in expression following long-term separation from a peer or from a mate.

**Figure 6.**
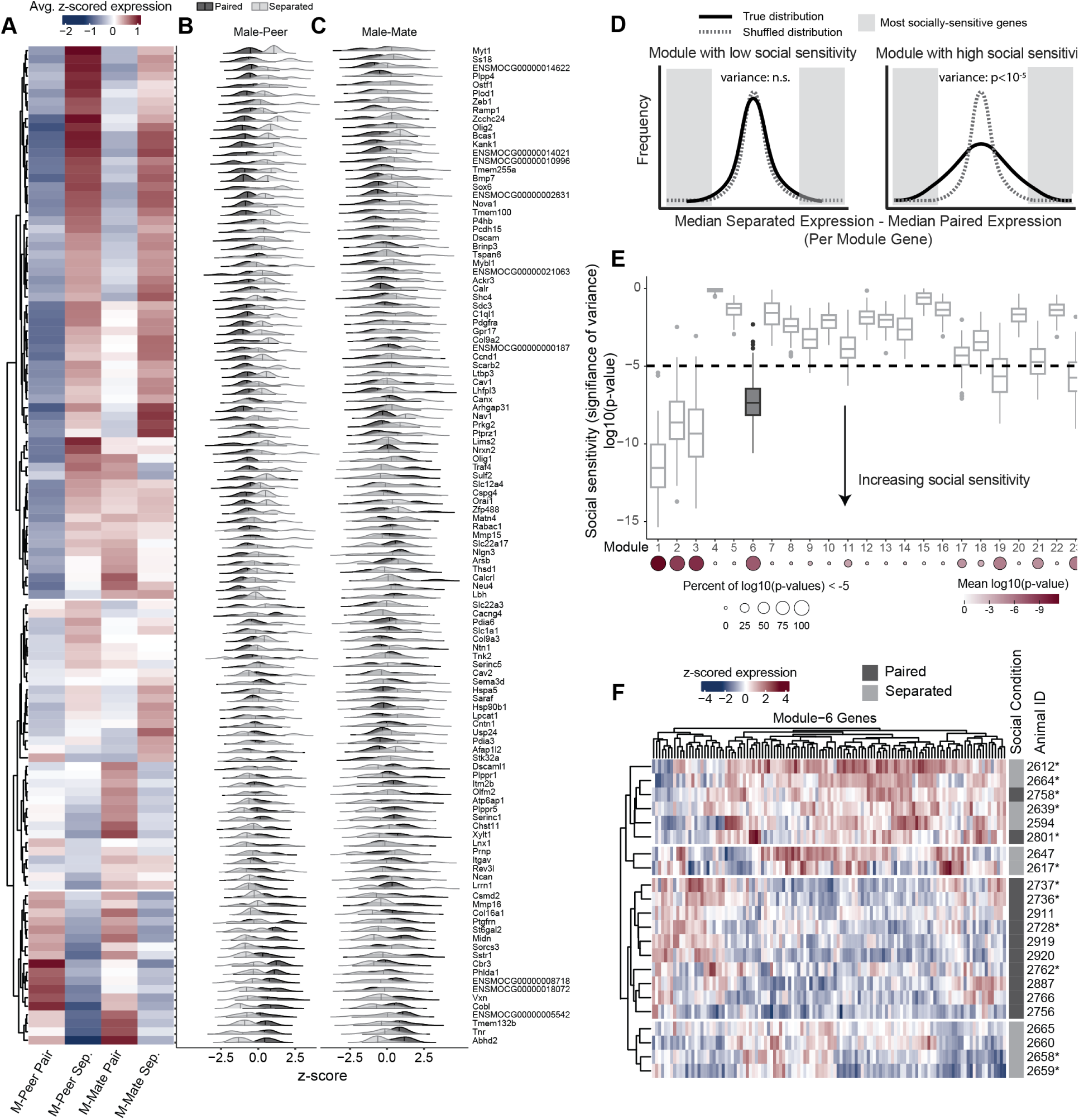
Module-6 genes are sensitive to bond loss. **A.** Heatmap of average z-scored normalized counts from RNA-seq data of Module-6 genes. Animals included were peer- and mate-paired males either paired for 2 weeks (paired) or paired for 2 weeks and then separated from their partner for 4 weeks (separated). **B, C.** Ridgeline plots of normalized counts on a per-animal basis for peer-paired (**B**) or mate-paired (**C**) animals. **D.** Schematic of difference score distribution comparisons. Modules with high social sensitivity will have many genes with large-magnitude differences between paired and separated groups, resulting in significantly more variance than expected by chance (shuffled). **E.** Social sensitivity of genes in each module. log10(p-value) distributions for F-tests comparing the variance of true (separated minus paired) difference scores vs. shuffled by animal ID (separated minus paired) difference scores for each module. Bottom: Dotplot with size representing the percentage of log10(p-values) < −5 and the color representing the mean(log10(p-value)) for each module. **F.** Heatmap of animal clustering based on Module-6 genes. Animals cluster separately by social condition but not by pair type within condition. *mate-paired.

We next asked whether Module-6 was more sensitive to partner separation than our other Hotspot gene modules. For each module gene, we found the difference in median expression level between paired animals and separated animals (combining peer and mate-paired groups, z-scored expression per-animal) (Fig 6B, C). This generated a distribution of difference scores for each module. If module genes are particularly socially sensitive, their true distribution of differences should have a high variance, representative of large differences between paired and separated animals. To determine the statistical significance of variance for each module, we iteratively shuffled the animal identities prior to calculating the same metrics to generate a null distribution of difference scores (schematic example provided in Fig 6D). Using an F-test, we tested whether the true module difference score distribution was more variable than chance for each iterative shuffle, repeated 100 times. We found that Module-6 had significantly greater variance than the shuffled null distributions (p = 10^−7.19^; Fig 6E), and 91% of the shuffled iterations for Module-6 had p-values smaller than a cutoff of p ≤ 10^−5^. Module-6 showed more significant variance than all but 3 other modules, indicating that Module-6 genes are highly sensitive to bond loss and to social environment more generally (paired vs separated). Interestingly, the other 3 modules that show the greatest variance (social sensitivity) are all enriched in glial clusters (Module-1 in microglia, Module-2 in astrocytes, and Module-3 in mature oligodendrocytes, Fig 2E), suggesting that several glial cell types are sensitive to partner separation.

Finally, we demonstrated that paired and separated animals could be distinguished by their transcriptional profiles of Module-6 genes. We used hierarchical clustering to group individual animals based on their expression of Module-6 genes. We found that animals largely clustered by whether they remained paired or were separated from their partner, irrespective of whether they were in peer or mate bonds (Fig 6F). This indicates that Module-6 genes are socially regulated but not sensitive to relationship type, consistent with our observation that transcriptional synchrony does not differ based on relationship type (peer or mate).

Together, our data indicate that Module-6 and OPCs are sensitive to bond loss at the level of individual pairs. This suggests a model in which Module-6 genes converge with a partner when a bond is formed, and that this pair-specific transcriptional signature erodes upon partner separation. This is consistent with prior conclusions from this dataset (*23*) and provides further evidence that oligodendrocytes may play an important, yet understudied, role in social bonds.

## Discussion

Using single nucleus transcriptomics in prairie voles, we provide new insights into the cellular basis of peer and mating-based social bonds. Our data indicate that different gene programs correlate with affiliative behavior in different sexes and relationship types, and that pairs exhibit similarity in cellular transcriptional states regardless of relationship type, which we refer to as transcriptional synchrony. Further, we show that a gene module expressed in OPCs is synchronized between partners and sensitive to partner loss. Our data suggest that a pair-specific transcriptional state supersedes a sex-specific or pair type-specific transcriptional state, and that the degree of intra-pair transcriptional similarity may influence, and be influenced by, pairwise behavior. Thus, transcriptional synchrony may serve as a pair-specific mechanism that contributes to the organization of successful intra-pair behavior.

One intriguing finding from our paper is the potential importance of oligodendrocytes in transcriptional synchrony and pair bonding behavior, confirming prior transcriptional work that likewise suggested a role for oligodendrogenesis in pair bonding (*23*). Social bonds are underpinned by a form of complex social learning as partners develop a repertoire of shared behaviors. Work in mice has shown that myelination is important for encoding and stabilizing complex memories (*26–28*). Myelination in the mammalian brain is also sensitive to social environment (*36*, *37*). This raises the intriguing but speculative possibility that oligodendrocytes contribute to bonding via myelination of neurons that encode key facets of behavior.

Overall patterns of transcription rather than expression of any single gene likely contribute to the emergence of a shared, bonded neural state. However, we did note that Down syndrome cell adhesion molecule (*Dscam*) is among the synchronized Module-6 OPC genes. *Dscam* is important for synaptogenesis, neuronal patterning, and is a risk gene for autism spectrum disorders; its disruption in mice leads to social behavior and learning deficits (*38–40*). *Dscam* is also one of 24 genes whose differential expression is conserved in taxonomically diverse monogamous species (*41*). Thus at least one gene in Module-6 is functionally consistent with genes that are likely to guide social behavior and bonding.

In the brain, neuronal firing and transcription mutually affect one another, providing a mechanism which may link neural synchrony and transcriptional synchrony. This is corroborated by evidence that expression of the immediate early gene *cFos* becomes correlated between prairie vole partners within the first day of pairing, likely contributing to parallel downstream effects on transcription. In addition to neurons, neural activity sculpts transcriptional profiles in glia (*42–47*), and in oligodendrocytes specifically, neural activity affects proliferation, differentiation, and myelination, which in turn modulates neural activity (*48*). Thus, it is possible that shared neural activity patterns during social interactions could have wide-ranging effects on transcription in oligodendrocytes (or other cells) in two individuals. Ultimately, neural activity-transcription relationships may be mutually reinforcing, with neural activity driving the initiation of transcription, which in turn creates a shared transcription state that is likely to lead to more aligned neural activity. Strengthening of this over the course of a relationship may contribute to organized behavior and other pairwise facets of bonding.

The factors that drive shared transcriptional state and/or those that contribute to variation in synchrony across pairs remain unknown. Results from our transcriptional decoder suggest that no single factor (cohort, sex, relationship type) was sufficient to drive the same degree of synchrony between animals observed in bonded pairs. Thus, we postulate that shared socio-environmental factors, including ongoing navigation of social relationships, is a major driver of transcriptional synchrony. Prior work also suggests that at least some aspects of transcriptional synchrony can emerge within the first few hours of interaction; in fighting betta fish, brain-wide transcriptomes become correlated within 60 minutes of interaction, and in prairie voles, *cFos* expression becomes correlated within 24 hours. This raises the intriguing possibility that the most salient and dynamic features of an environment – namely social interactions with a cohoused partner – are a primary and heterogeneous driver of transcriptional synchrony. Here, we build on this by further showing that transcriptional synchrony is also evident at longer timepoints, potentially helping to cement bonds over time.

While we provide compelling evidence of transcriptional synchrony, our studies have a number of limitations and important future directions. For instance, the specific factors that contribute to variation in synchrony *across pairs* remain unknown. Likewise, the functional relevance of transcriptional synchrony has yet to be examined. To what extent does transcriptional synchrony depend on affiliative bonds versus other types of social interactions? Finally, it will be important to determine whether this phenomenon is observed across brain regions or even across tissues.

In sum, the strengthening of transcriptional synchrony may facilitate a pair’s ability to coordinate their behavior. Regardless of the specific drivers of this phenomenon, similar transcriptional profiles may provide a common biological substrate that facilitates enhanced neural synchrony and organized behavior. This alignment of transcription and neural activity may enable partners in long-term, meaningful relationships to predict the actions of their partner, quickly infer their mental state, and choose an appropriate behavioral response.

## Methods

### Animals

Prairie voles were bred in-house in a colony descended from wild animals collected in Illinois, USA. Animals were weaned at 21 days and housed in same-sex groups of 2-4 animals in standard static rodent cages (19l × 10.5w x 5 in. h) with ad-libitum access to rabbit chow (5326-3, PMI Lab Diet) and water. Sunflower seeds and dried fruit treats were given to supplement the diet, and cotton nestlets, PVC tubes, and plastic houses were given for enrichment. All voles were between 77-154 days old at the start of the experiment. On day one of the experiment, animals were co-housed with either a same-sex (peer-paired) or opposite-sex (mate-paired) animal in smaller standard static rodent cages (11l x 8w x 6.5h) with ad-libitum access to food and water, cotton nestlets and plastic houses. These pairings resulted in 11 female/male pairs, 4 female/female pairs, and 5 male/male pairs. The colony room was kept at 23-26C with a 14:10 light:dark cycle. All procedures were performed during the light phase and approved by the University of Colorado Institutional Animal Care and Use Committee.

### Partner preference test

Partner preference tests were performed on day 14 of the experiment. Both partners within a pair were tested consecutively, with the order randomized. Briefly, partner preference tests were performed in a three-chamber plexiglass arena (76 cm long x 20 cm wide x 30 cm tall). Opaque dividers were placed between the chambers of the arena to prevent animals from seeing each other at the start of the test. Partner and novel animals were tethered to opposite ends of the chamber using tethers consisting of a wall-attached eye bolt with a chain of fishing swivels. Animals were briefly anesthetized with isoflurane and a zip tie was used around the animal’s neck to attach them to the tether. Rabbit chow and water were given for the duration of the test. At the start of the test, the experimental animal was placed in the center chamber and the dividers were removed, allowing the experimental animal to freely roam the arena for 2.5 hours. The tests were recorded using overhead cameras (Panasonic WVCP304).

The videos were analyzed using TopScan software v3.0.0 using the parameters in Ahern et al (*49*). Behavior was analyzed using a custom Python script developed in-house (https://github.com/donaldsonlab/CleversysSummaryRearranger) to calculate time spent huddling with the partner and time spent huddling with the novel. Partner and novel proximity metrics consisted of time spent in the chamber with the respective animal, respectively. Partner preference was calculated as (partner huddle time/[partner huddle time + novel huddle time]) *100%.

### Free interaction test

Free interaction tests were performed on day 15 of the experiment as described in Brusman et al (*33*). Each pair was placed in a plexiglass arena (50.7 cm long x 20 cm wide x 30 cm tall) and allowed to freely roam the arena for 3 hours. The tests were recorded using overhead cameras (Logitech C925e webcam). The videos were scored for total interaction time post hoc using TopScan software v3.0.0 and the parameters described in Brusman et al.

### Tissue harvest

On day 16 of the experiment, animals were rapidly decapitated using a guillotine and brains were placed immediately on dry ice before moving to −80C. Uteri of female animals were removed and dissected to assess pregnancy status. Brains were later sectioned on a cryostat until the front of the NAc was reached (coordinates). A 2 mm diameter tissue punch was used to punch the NAc ∼1mm deep on each side of the brain. Punches were stored separately for each animal in eppendorf tubes at −80C until nuclei dissociation.

In our experiment, of our 11 mate-paired females, 8 became pregnant over the course of pairing, while 3 did not. While it is possible that pregnancy could be a confounding factor in behavior or the transcriptional patterns we observed, we found no differences in partner huddle time between pregnant and non-pregnant females (Wilcoxon rank-sum test, p = 0.92), and a difference in gene expression only in Module-19 (Wilcoxon rank-sum test, p = 0.017). Of note, Module-19 is not a synchronized module, nor is it a module that correlates with PPT behavior. Additionally, because there are 8 pregnant and 3 non-pregnant animals in this sample, this statistical difference may be confounded by sample size.

### Single nuclei isolation

Brain punches were placed on wet ice until barely thawed. 0.5 mL Complete Buffer HB (250mM sucrose, 25mM KCl, 5mM MgCl2, 20mM Tricine-KOH, 0.042% BSA, 0.06 U/ul RNAsin, 0.15mM spermine, 0.5mM spermidine, 1mM DTT, cOmplete EDTA-free protease inhibitor cocktail tablet) was then added to each tube with brain punches and then Complete Buffer HB and punches were transferred to 2 mL dounce homogenizer. Samples were dounced with a tight pestle 20 times. 16 uL 5% IGEPAL CA-630 in Buffer HB (250mM sucrose, 25mM KCl, 5mM MgCl2, 20mM Tricine-KOH) was added to the dounce homogenizer and the samples were dounced 10 more times. The samples were then filtered through a Flowmi 40 um strainer into a 2 mL LoBind Eppendorf tube. The volume of flowthrough was measured and an equal volume of Working Solution (50% iodixanol (OptiPrep), 25mM KCl, 5mM MgCl2, 20mM Tricine-KOH, 0.042% BSA, 0.064 U/ul RNAsin) was added to the homogenate and mixed by gentle pipetting using a wide-bore P1000 pipette tip. First, 50% iodixanol was made by diluting 60% iodixanol (OptiPrep) in Working Solution. Then, 30% and 40% iodixanol solutions were made by diluting the 50% iodixanol/Working Solution in Complete Buffer HB. In 0.8 uL ultracentrifuge tubes, 160 uL 30% iodixanol was carefully layered on top of 80 uL 40% iodixanol and the interface was labeled with pen. Approximately 560 ulof filtered sample was carefully layered on top of the iodixanol gradient using a 23G angled needle attached to a 1 mL insulin syringe. The samples were centrifuged at 10,000g for 30 min in a swinging bucket ultracentrifuge at 4C with the acceleration set to slow and the brake set to slow. When the spin was finished, the majority of the layer above the interface was carefully removed using a 23G angled needle attached to a 1 mL insulin syringe. Approximately 70 uL of nuclei were collected at the 30%-40% iodixanol interface using a 14G blunt end needle attached to a 1 mL insulin syringe and put in a LoBind eppendorf tube. To check the concentration of nuclei, 10 uL trypan blue was added to 10 uL nuclei and nuclei were examined using a brightfield microscope at 10X magnification. Samples were diluted to 320-2020 nuclei/uL in Complete Buffer HB before loading onto 10X Genomics Chromium controller.

### Sequencing

Samples were sequenced using the Chromium next GEM Single Cell 5’ kit v2 from 10X Genomics. Single nuclei suspensions were loaded onto the 10X Genomics Chromium controller at concentrations between 320-2,020 nuclei/ul, aiming to capture 5,000 nuclei/sample at a read depth of 40,000 reads/nucleus. Single nuclei RNA-seq (snRNA-seq) libraries were prepared according to the manufacturer’s instructions. Library quality was assessed using the Agilent High Sensitivity D5000 ScreenTape System and were subsequently sequenced using paired-end sequencing on an Illumina NovaSeq6000.

### Analysis

#### Sequence alignment and UMI counting

Raw reads were aligned and counted using the Cellranger v3.1.0 analysis pipeline. Briefly, a Cellranger-compatible genome was generated via the mkref function using the published prairie vole genome (MicOch1.0, GenBank accession number AHZW00000000). Cellranger mkfastq was then used to create FASTQ files from the raw BCL files. Across samples, we had a mean percent read alignment of 80.3%. Cellranger count was then used to align, filter, and count the reads. The mean number of nuclei per sample was 4,862 and the mean read depth was 37,745 reads/nucleus. Samples with a mean less than 20,000 reads/nucleus were excluded from downstream analysis (1/40 samples).

#### Sample integration and clustering

We used Seurat v4.3.0 in R v4.2.2 to analyze our single nuclei data (*50*). First, the filtered feature barcode matrices were used to create Seurat objects with sample-associated metadata. For each sample, cells were filtered to have >200 genes detected per cell and <5% reads from mitochondrial genes. The 39 samples were then transformed using Seurat’s SCTransform function and integrated using Seurat’s SCT Integration. This left 178,020 nuclei. Following integration, potential effects of the sequencing cohort and the behavior cohort were regressed out. Cells were re-clustered, and clusters were removed that expressed *Slc17a7* (excitatory neurons) or *Pecam1* (endothelial cells). These are two molecular markers consistent with cell types adjacent to, but not found within, the NAc (*51*). Cells were clustered again, and one remaining cluster was removed that expressed *Trh* (hypothalamus). Finally, this yielded 142,488 nuclei included in the final clustering analysis.

Using the SCT-normalized counts, dimensionality reduction was performed using PCA. The top 50 PCs were used for UMAP clustering to generate the final cell clusters. 15 clusters were identified and their biological identities were determined by the expression of known marker genes for cell types in the NAc.

We then calculated the percentage of cells in each cluster for each animal. To compare groups, we created generalized linear mixed models with sex and pairing type as fixed effects using the glmmTMB package (v1.1.8) in R (v4.2.2). For post-hoc comparisons, we used the emmeans package (v1.8.5) in R with p < 0.05 set as our significance threshold for all experiments.

#### Hotspot analysis

We used the Hotspot (v1.1.1) analysis pipeline (*24*) in Python (v3.11.3) to group genes into gene modules based cross-correlations in gene expression across cells. First, we converted our Seurat object to a SingleCellExperiment object. Next, we created the Hotspot object using the Seurat SCT counts matrix, the PCA previously calculated by Seurat, and a negative binomial model. We then computed the neighborhood using an unweighted graph and 30 nearest neighbors. We computed the autocorrelations for each gene to determine the top 3000 most variable genes, and then computed pairwise local correlations between these genes. Finally, we created our modules using these local correlations, with a minimum module size of 30 genes. This left us with 23 modules ranging in size from 31-415 genes. We then calculated module scores for each module in each cell to determine module enrichment across cell populations.

Downstream of the module creation, we calculated the average (mean) gene expression for all genes in each module across all cells for each animal. To determine group differences, we generated generalized linear mixed models using glmmTMB (v1.1.8) with sex and pairing type as fixed effects and did post-hoc testing using emmeans (v1.8.5) as stated above.

To determine whether partners were more similar than non-partners, we began by calculating a Spearman correlation for the average module gene expression between partners. We then used the SCT counts for the genes in each module to find the distance between partners in Euclidean space. As a control, we found the distance between all non-true pairs (chance) and compared the true partner distances to that distribution. Finally, for each module, we ranked the animals according to their average module gene expression and found the rank-based distance between partners. With 38 paired animals, the expected rank distance between any two animals by chance is 13. We performed one-sample t-tests for each module against a null value of 13.

To determine the roles of the genes in each module, we performed gene ontology (GO) analysis using the gprofiler2 package (v0.2.1) in R. The prairie vole gene names were mapped onto the ENSG namespace prior to running GO analysis via gprofiler2. All GO p-values were FDR corrected.

#### Support vector machine analysis

We used the e1071 package (v1.7-13) in R (v4.2.2) to analyze our data using support vector machines (SVMs). For each cluster, we downsampled our cells to 200 cells per animal if there were >200 cells for that animal in that cluster. For our SVMs that were able to classify the animal as itself, we split the cells into a training set and a test set, where 75% of the cells were randomly allocated to the training set and the remaining 25% to the test set. We then trained the SVM for each cluster to classify cells based on animal identity using the scaled Pearson residuals of the gene expression of the top 3,000 variable genes as determined by Seurat. For each animal, we found the percentage of cells from the test set that were classified as the animal from which they originated (self), the partner of the animal from which they originated (partner), or any other animal (other). We then compared the distributions of “self”, “partner”, and “other” classifications.

To train our SVMs that were not provided with the option to classify as self, we downsampled our cells as mentioned above, and then trained a separate SVM for each animal for each cluster. For each SVM, we held a single animal out of the training set, and then tested the SVM on only the held-out animal. This forced the SVM to classify each cell as an animal other than the animal from which it originated. We then calculated the percentage of cells in the test set classified as the partner or as other and compared these groups using generalized linear mixed models with estimated marginal means as post-hoc tests.

#### Partner separation experiment

The partner separation experiment was performed as described in Sadino et al (*23*). Briefly, the animals from Sadino et al. included in this analysis were sexually naïve male prairie voles paired with either a same-sex (peer/sibling) or opposite sex (mate) partner for 2 weeks. On day 14, all experimental animals underwent a PPT. The “paired” voles were then cohoused for an additional two days (16 days total, consistent with snRNA-seq experimental animals) and the “separated” voles were separated from their partner for 4 weeks. At these final timepoints, animals were sacrificed by rapid decapitation and the NAc was dissected out from the brain prior to processing for RNA-seq.

#### Analysis of RNA-seq separation data

Sequence mapping and counting was performed as described in Sadino et al (*23*). Following this analysis, DESeq2 (v1.38.3) (*52*) was used in R (v4.2.2) to calculate the normalized counts on a per-animal basis. For each gene, the normalized counts were z-scored. For the Module-6 analysis specifically, we found that 117/143 Module-6 genes were detected in the RNA-seq dataset. For each of these 117 genes, we calculated the mean z-score within groups and used hierarchical clustering to cluster the genes into 3 primary clusters. We then used the z-scored counts on a per-animal basis to cluster individual animals. The animals largely separated by social condition (paired vs. separated).

Finally, we combined the mate-paired and peer-paired animals into two groups based on social condition: paired and separated. For each group in each module, we found the median expression for each gene. We then used those median values and calculated the difference between groups for each gene as a metric representing the degree of change between paired and separated animals. This resulted in a distribution of change scores for each module. As a control, we iteratively shuffled the animal identities 100 times prior to performing these same calculations, and then used an F-test to compare the two distributions for each iteration.

#### Code availability

All code written for this analysis is provided on the Donaldson Lab Github at: https://github.com/donaldsonlab/snRNAseq_voles.

## Acknowledgments

We thank Jay Hesselberth and his lab for allowing us to use their space to perform the nuclei isolations. We also thank the Genomics Core at CU Anschutz for sequencing our samples. We thank Jessica Abazaris and the rest of the animal care staff at the University of Colorado Boulder for their excellent care of the voles, and the voles themselves for their sacrifice. We thank Zachary Johnson for discussing single nucleus data, and Catherine Peña and Yevgenia Kozorovitskiy for feedback on initial drafts of the manuscript.

## Funding

National Institutes of Health grant 1F31MH132278-01A1 (LEB)

National Institutes of Health grant 5T32GM008759-20 (LEB)

National Institutes of Health grant 1T32GM142607-01 (LEB)

Whitehall Foundation Award (ZRD)

Dana Foundation Award (ZRD)

National Institutes of Health grant DP2MH119421 (ZRD)

National Institutes of Health grant UF1NS122124 (ZRD)

National Institutes of Health grant R01MH125423 (ZRD)

National Institutes of Health grant IOS-2045348 (ZRD)

National Institutes of Health grant U01NS131406 (ZRD)

National Institutes of Health grant R01HL156475 (RDD, MAA)

Anna and John J. Sie Foundation/Global Down Syndrome Foundation (MAA).

## Author contributions

Conceptualization: LEB, ZRD

Formal analysis: LEB, MAA

Funding acquisition: ZRD, LEB, MAA, RDD

Investigation: LEB, ACF

Project management: ZRD

Resources: ZRD

Software: LEB, MAA

Mentorship: ZRD, MAA

Visualization: LEB

Writing – original draft: LEB, ZRD

Writing – review & editing: LEB, ZRD, MAA, RDD

## Competing interests

The authors declare the following competing interests: MAA and RDD have a patent for measuring transcription factor activity from eRNA activity. RDD was a co-founder of Arpeggio Biosciences. All other authors have no competing interests.

## Data and materials availability

All sequencing data is deposited on GEO (GSE255620) and will be made available upon publication. All code generated for the analysis of this data will be available on the Donaldson Lab Github (https://github.com/donaldsonlab).

## Ethics

All animal procedures were carried out in accordance with standard ethical guidelines (National Institutes of Health Guide for the Care and Use of Laboratory Animals) and approved by the Institutional Animal Care and Use Committee (IACUC) at the University of Colorado Boulder.

## Notes

### Summary of Updates

Addition of figures 5 and 6.

