## Supplementary Figures for "Single nucleus RNA-sequencing reveals transcriptional synchrony across different relationships"

Science
AAAS

### Supplementary Materials for

#### Single nucleus RNA-sequencing reveals transcriptional synchrony across different relationships

**Authors:** Liza E. Brusman[1], Julie M. Sadino[1], Allison C. Fultz[2], Michael A. Kelberman[1], Robin D. Dowell[1,3], Mary A. Allen[1,3*†], Zoe R. Donaldson[1,2*†]

†These authors contributed equally

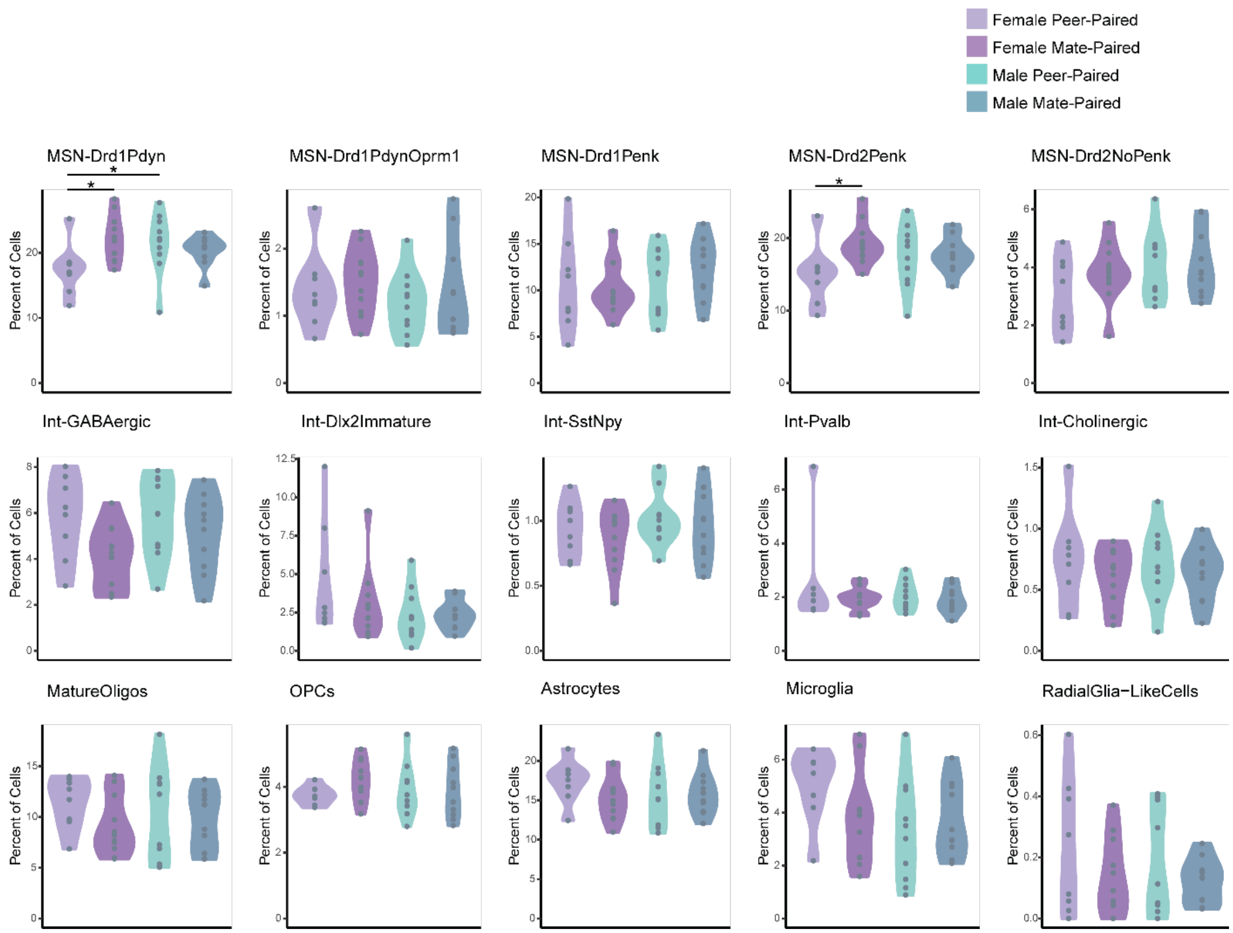


**Figure S1. Percentage of cells in each cluster by group.**

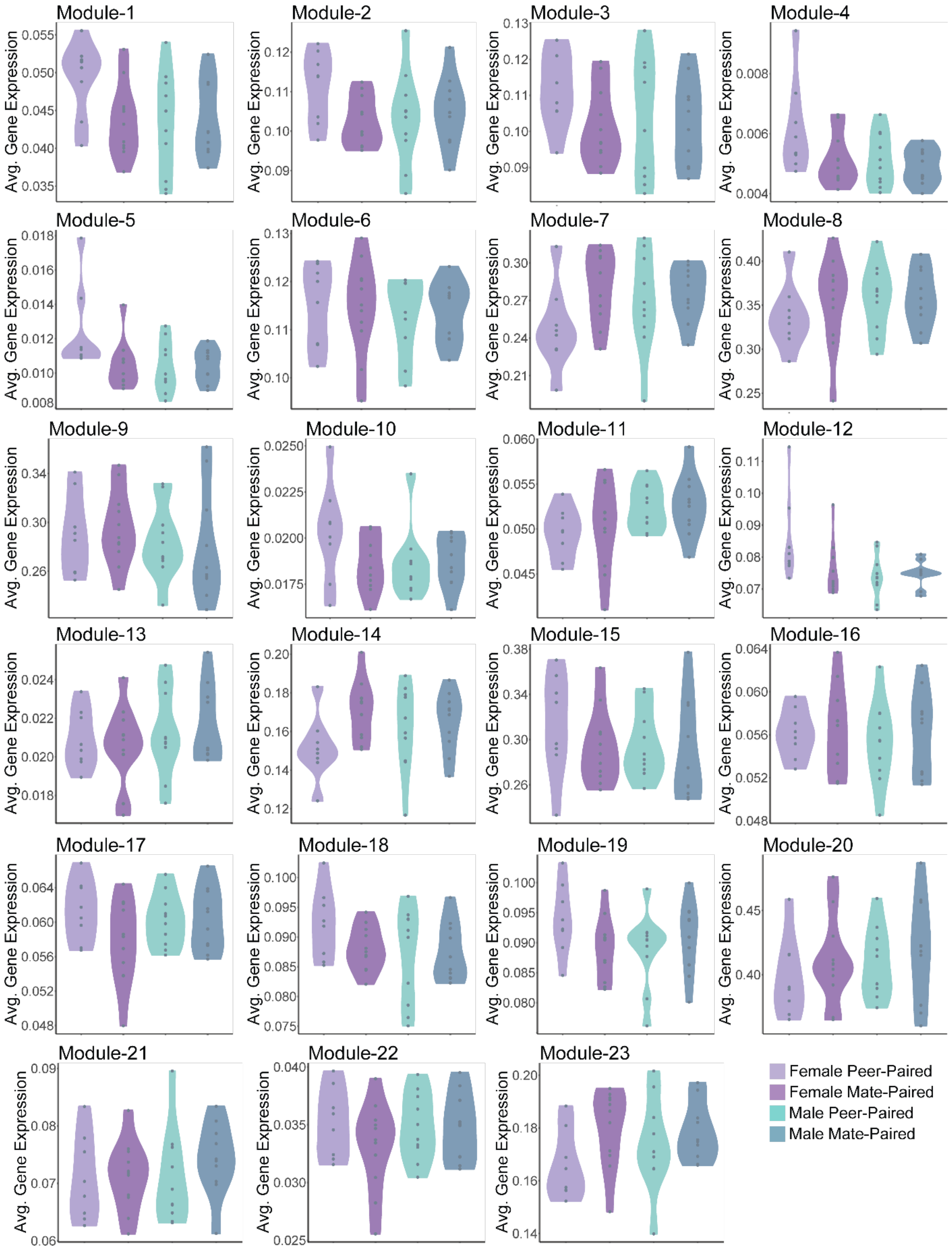


**Figure S2. Average module gene expression for all modules by group.**

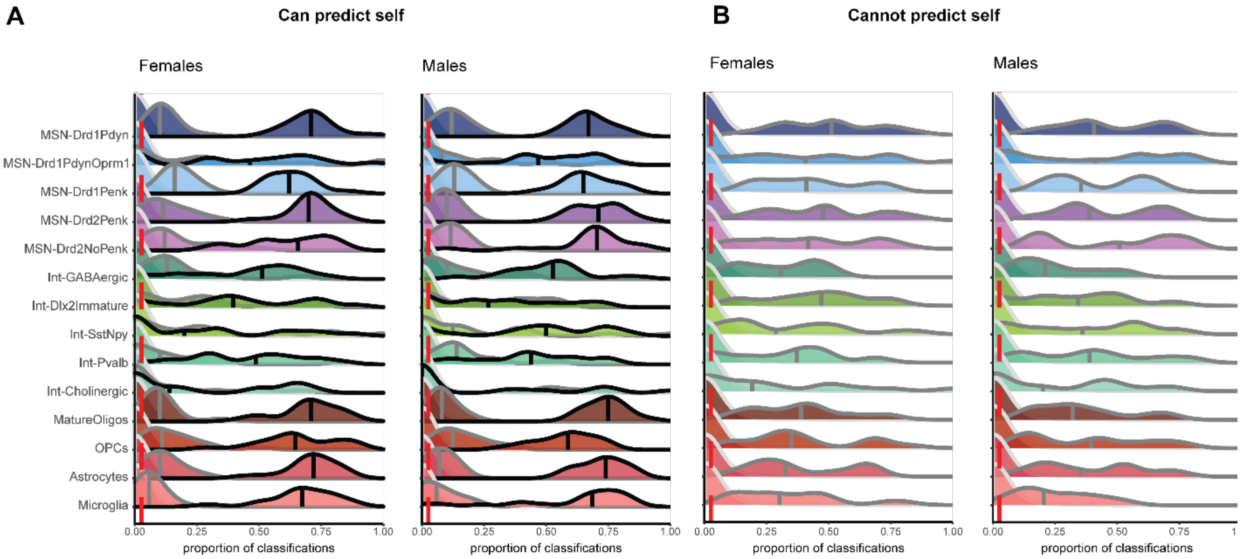


**Figure S3. SVM classifications by sex. A.** SVM classifications for SVMs that can predict "self." **B.** SVM classifications for hold-one-out SVMs that cannot predict "self."

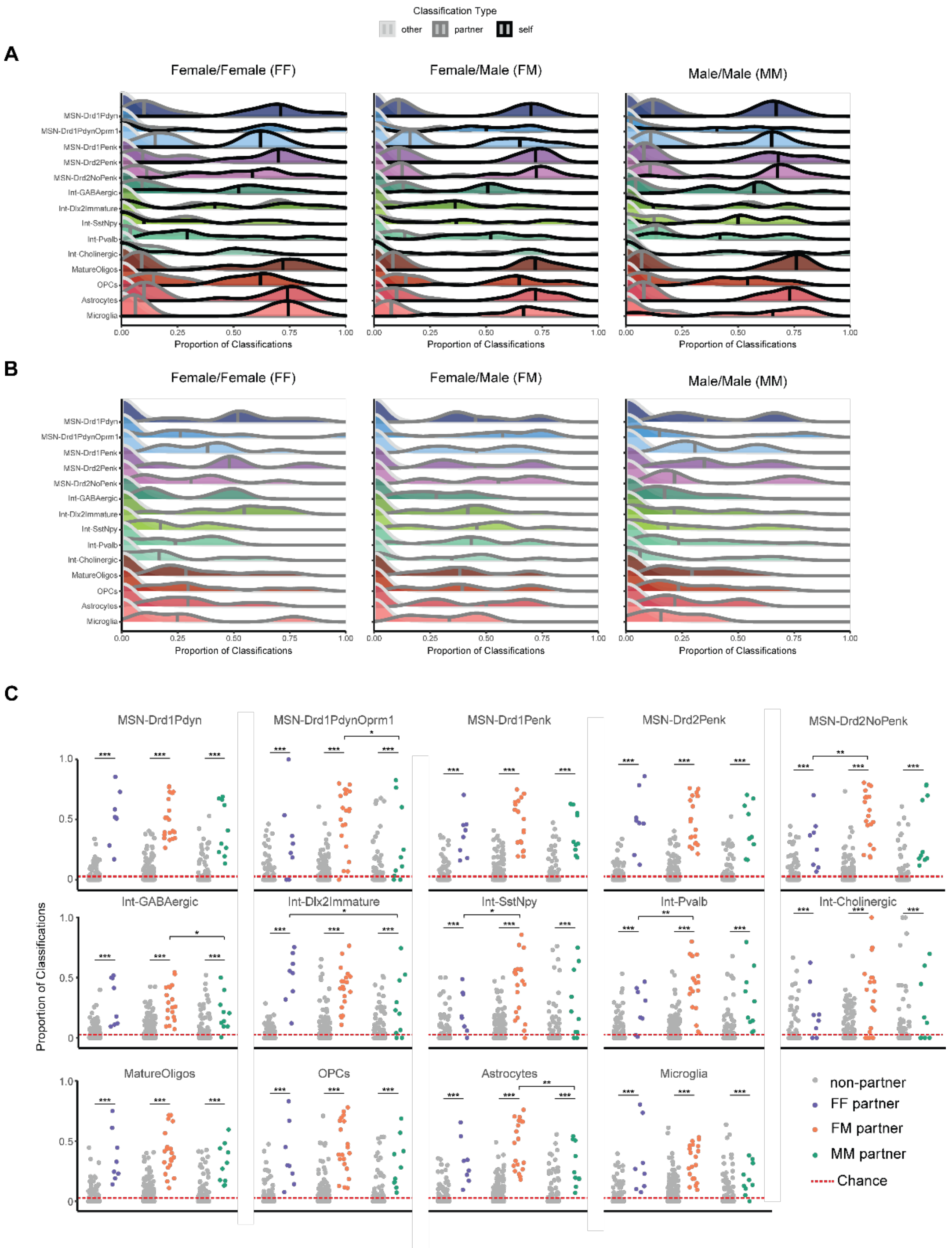


**Figure S4. SVM classifications by pairing type. A.** SVM classifications for SVMs that can predict "self." **B.** SVM classifications for hold-one-out SVMs that cannot predict "self." **C.** SVM classifications split by group compared to all "other" classifications on a per animal/per animal basis. Each point represents classifications of one animal for one other animal (e.g. proportion of classifications from Animal 1 cells as Animal 2 cells, Animal 3 cells, etc.). Line at y = 2.7% is chance probability of classification. * p < 0.05, ** p < 0.01, *** p < 0.001

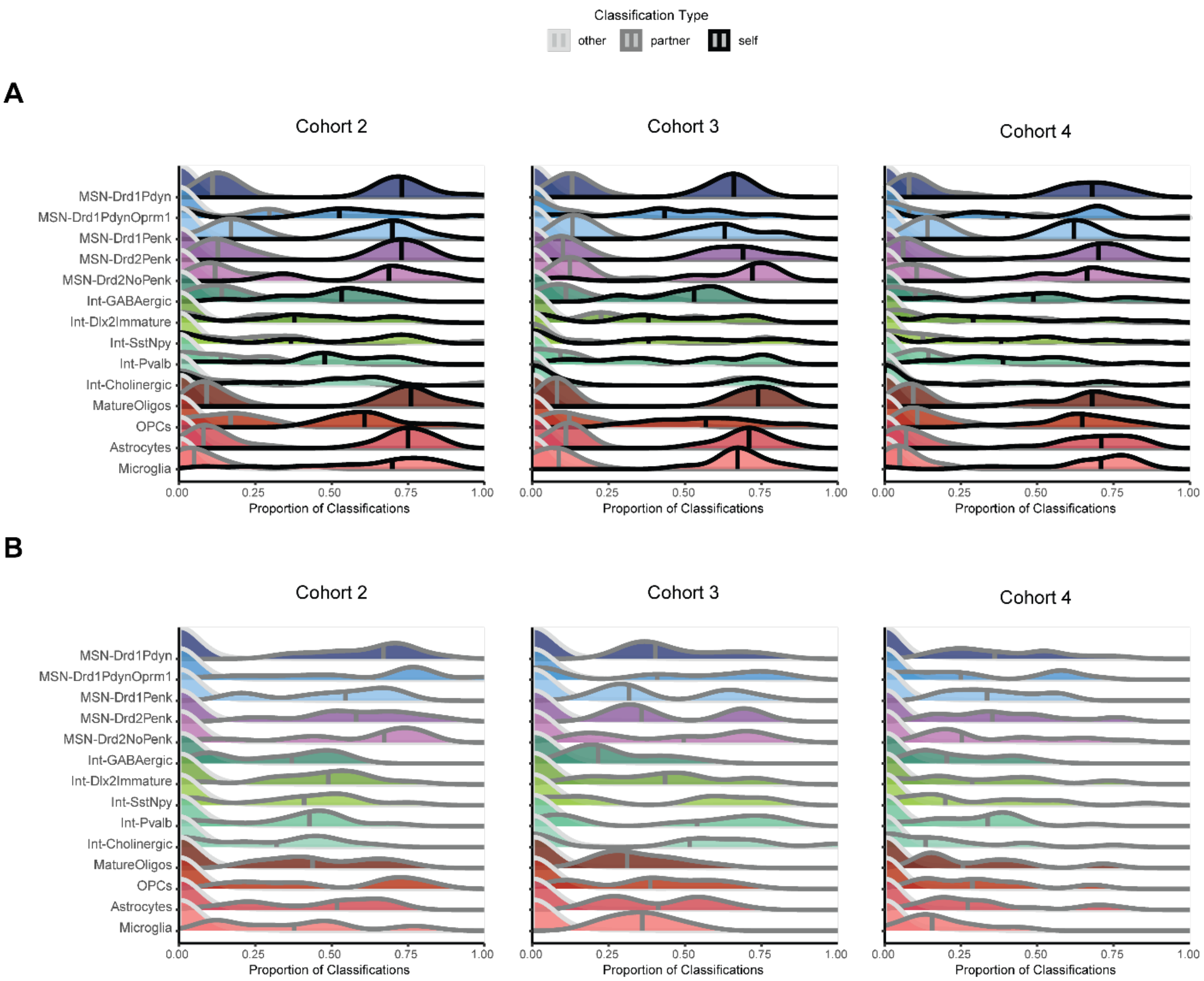


**Figure S5. SVM classifications by cohort. A.** SVM classifications for SVMs that can predict "self." **B.** SVM classifications for hold-one-out SVMs that cannot predict "self."

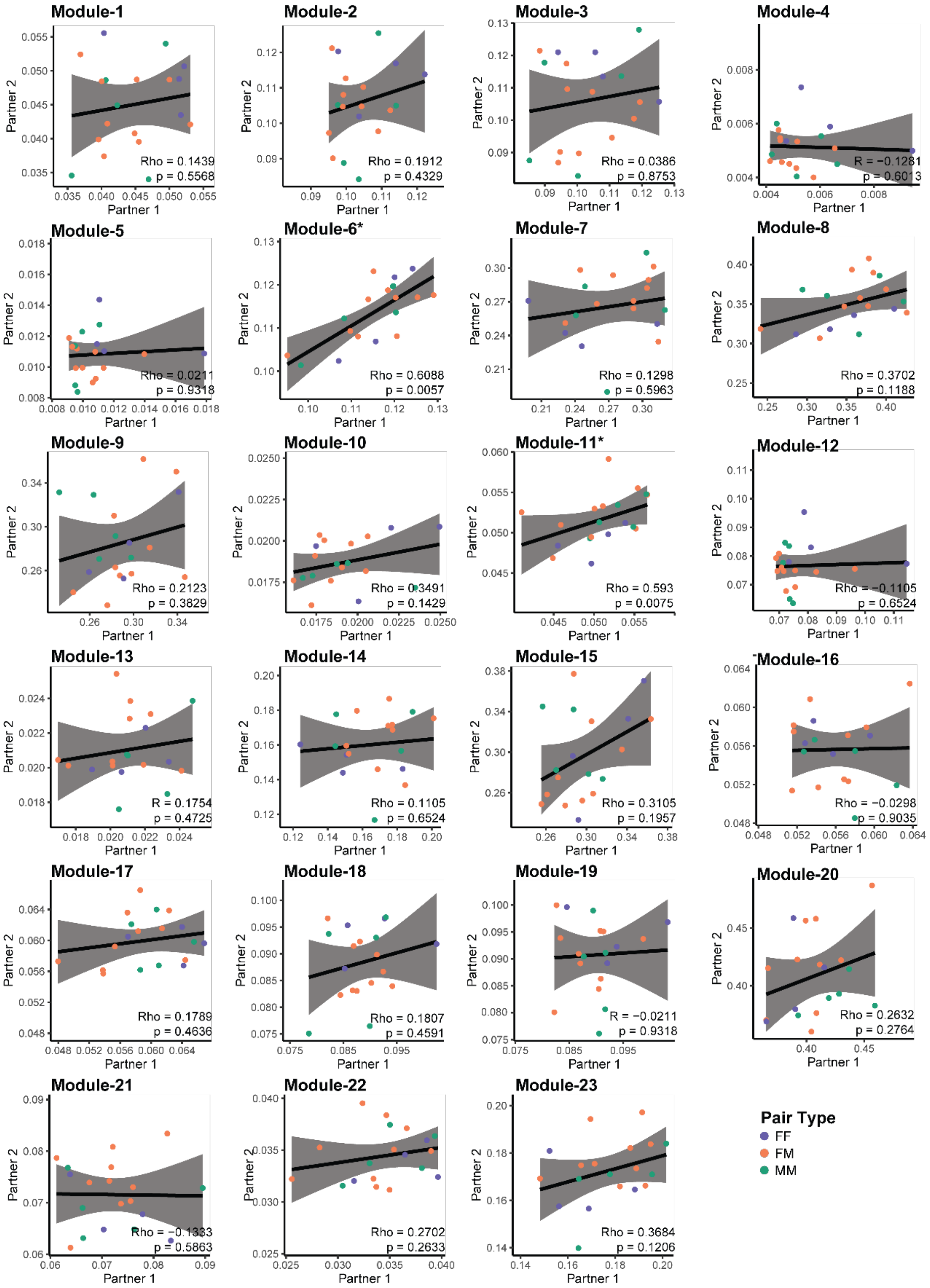


**Figure S6. Average module gene expression correlation between partners.** * $p < 0.05$

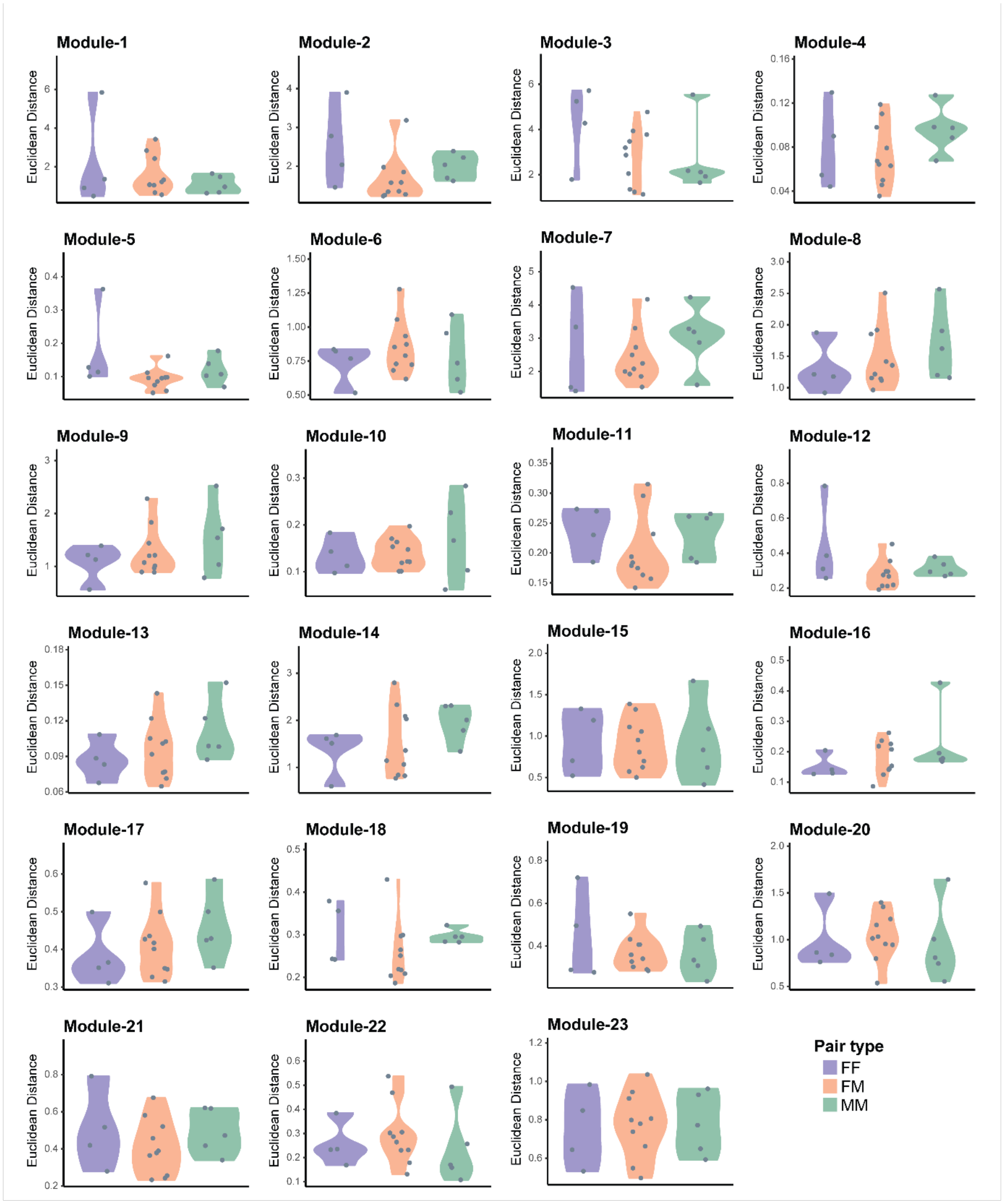


**Figure S7. Euclidean distance between partners for genes in each module, separated by pairing type.**

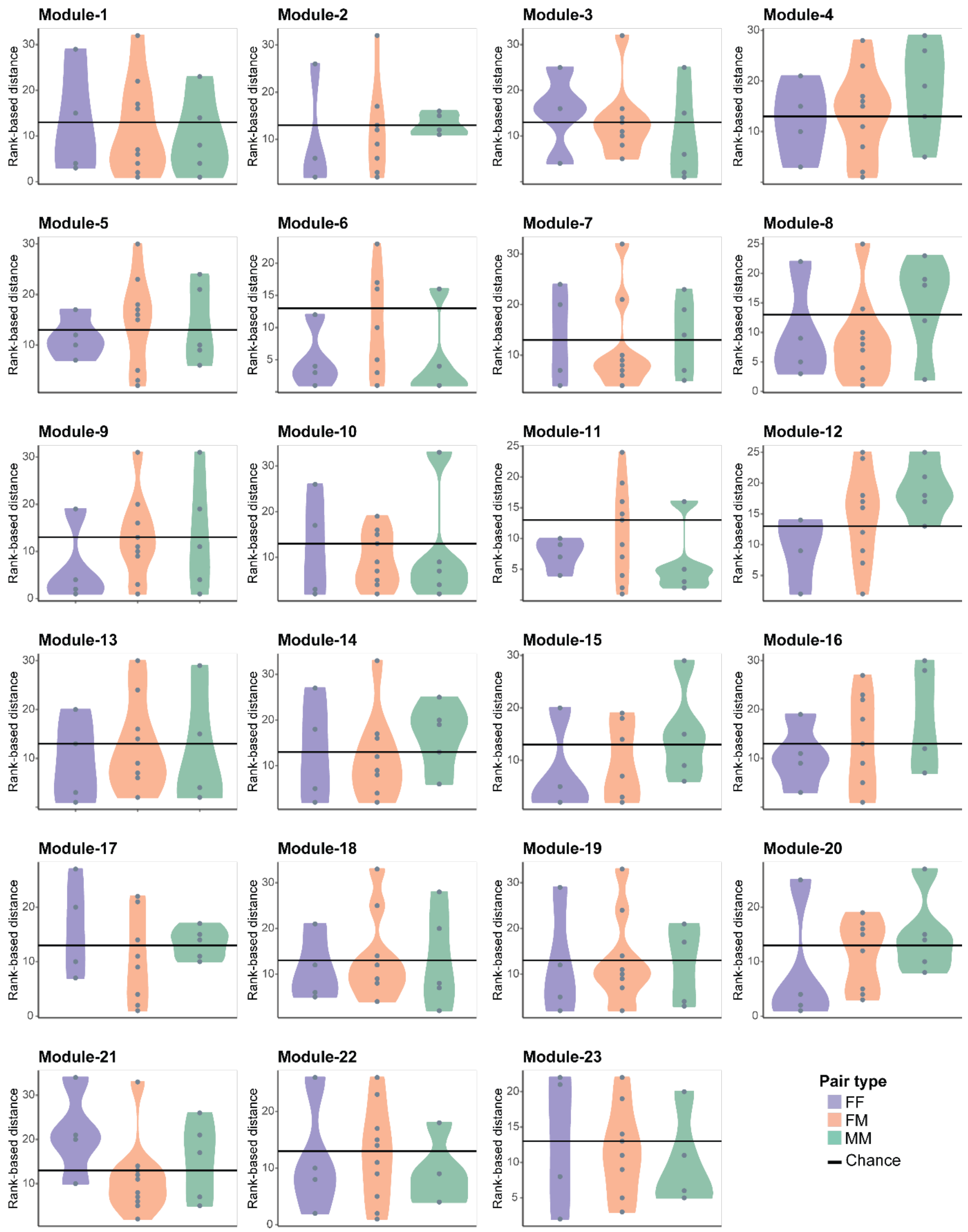


**Figure S8. Rank-based distances between partners separated by pairing type.**

**Table S1. (separate file)**
Sequencing metadata.
**Table S2. (separate file)**
Gene-module membership from Hotspot analysis.
**Video S1. (separate file)**
Video of the free interaction test.
